## Supplementary material for "Targeting Enhanced Digestibility: Prioritizing Low Pith Lignification to Complement low p-Coumaric Acid content as environmental stress intensity increase": supp figure S1

### Slide 1
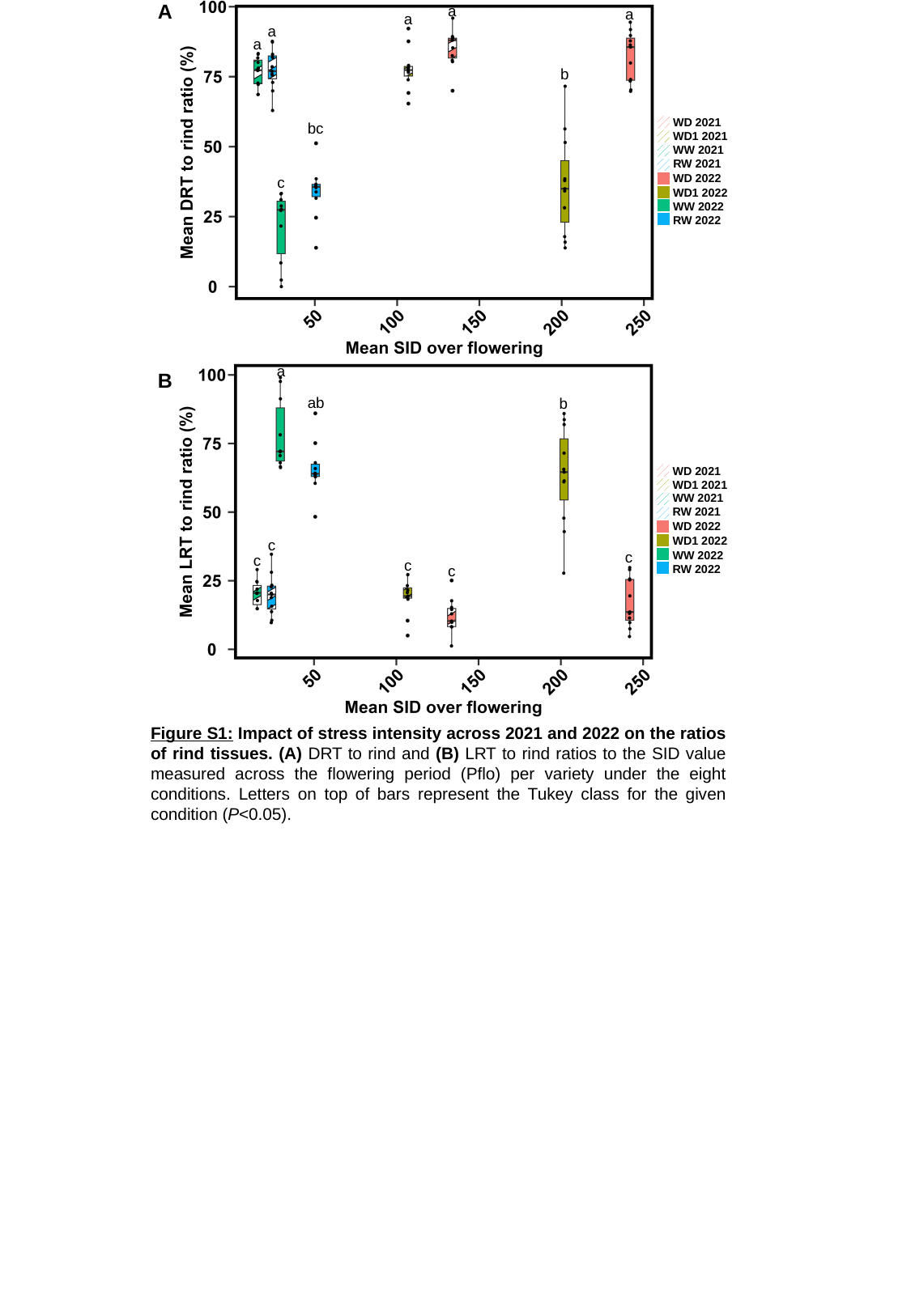

A
a
a
a
a
a
b
WD 2021
WD1 2021
WW 2021
RW 2021
WD 2022
WD1 2022
WW 2022
RW 2022
bc
c
a
B
ab
b
WD 2021
WD1 2021
WW 2021
RW 2021
WD 2022
WD1 2022
WW 2022
RW 2022
c
c
c
c
c
Figure S1: Impact of stress intensity across 2021 and 2022 on the ratios of rind tissues. (A) DRT to rind and (B) LRT to rind ratios to the SID value measured across the flowering period (Pflo) per variety under the eight conditions. Letters on top of bars represent the Tukey class for the given condition (P<0.05).
